# Exploring transient states of PAmKate to enable improved cryogenic single-molecule imaging

**DOI:** 10.1101/2024.04.24.590965

**Authors:** Davis Perez, Dara P. Dowlatshahi, Christopher A. Azaldegui, T. Bertie Ansell, Peter D. Dahlberg, W. E. Moerner

## Abstract

Super-resolved cryogenic correlative light and electron microscopy is a powerful approach which combines the single-molecule specificity and sensitivity of fluorescence imaging with the nano-scale resolution of cryogenic electron tomography. Key to this method is active control over the emissive state of fluorescent labels to ensure sufficient sparsity to localize individual emitters. Recent work has identified fluorescent proteins (FPs) which photoactivate or photoswitch efficiently at cryogenic temperatures, but long on-times due to reduced quantum yield of photobleaching remains a challenge for imaging structures with a high density of localizations. In this work, we explore the photophysical properties of the red photoactivatable FP PAmKate and identify a 2-color process leading to enhanced turn-off of active emitters, improving localization rate. Specifically, after excitation of ground state molecules, we find a transient state forms with a lifetime of ∼2 ms under cryogenic conditions which can be bleached by exposure to a second wavelength. We measure the response of the transient state to different wavelengths, demonstrate how this mechanism can be used to improve imaging, and provide a blueprint for study of other FPs at cryogenic temperatures.

## Introduction

Cryogenic electron tomography (cryoET) allows for the label-free observation of the organization and structure of molecules inside cells, frozen in their native-hydrated state. While the nanometer scale resolution is impressive, the contrast is too low to directly identify the majority of biomolecules, specifically for structures smaller than about 500 kDa, despite recent developments in computational methods.^1-3^ While reducing this size limitation through improved imaging and computational methods is an active area of research with great potential,^4^ there is also interest in developing labelling strategies for cryoET.^5, 6^ Cryogenic correlative light and electron microscopy (cryoCLEM) with standard fluorescence microscopy has often been used to identify the location of molecules of interest in tomographic reconstructions.^7-9^ CryoCLEM has the advantage of the wide library of fluorescent labels and labeling strategies developed for room temperature (RT) imaging, but the few 100 nm resolution obtainable with diffraction-limited light microscopy is two orders of magnitude worse than cryoET. To bridge this gap, super-resolution fluorescence microscopy techniques have been adapted to cryogenic temperatures (CT) and paired with cryoET, for a method called srCryoCLEM. Various super-resolution methods have been used, including single-molecule localization microscopy (SMLM)-based techniques such as cryo photoactivation localization microscopy (cryo-PALM) which are appealing as they have provided the highest localization precision to date (∼10 nm) in these cryoCLEM workflows.^10-13^

To adapt SMLM to CT, several key challenges needed to be overcome. SMLM requires a sparse subset of active emitters in each frame, which means emitters must turn on and off efficiently. At room temperature, a variety of active control mechanisms have been developed and optimized.^14, 15^ However, many photoswitching pathways accessible at RT occur with extremely low probability at CT. Previous work has identified a handful of FPs which photoswitch efficiently at 77K, including PAGFP, rsEGFP2, and PAmKate,^10, 16-18^ which facilitated the first generation of srCryoCLEM experiments. PAmKate absorption is shifted to long wavelengths (orange), which is useful to avoid cellular autofluorescence that occurs with blue or green pumping,^19^ and the protein starts in a dark state reducing the need for photobleaching or shelving before imaging. While molecules can be turned on efficiently, they remain on for extremely long times at CT. Therefore, these implementations were limited by the number of localizations which could be acquired in a given experimental time. The large number of photons collected allows for precise localization, but once a molecule’s position is known to a high enough precision, it needs to be turned off so the next molecule in a diffraction limited area can be localized. These long on times are caused by two factors: limited excitation intensity to avoid sample heating and reduced quantum yield of photobleaching.

For samples imaged in fluorescence to be compatible with subsequent cryoET, the ice must remain vitreous or amorphous throughout the whole experiment. This means the sample temperature must be held below 135 K,^20^ limiting the laser intensity which can be used to excite the fluorophores. Recent work found that the primary source of heating is absorption of excitation light by the substrate, which is traditionally a thin carbon film.^21^ Because thermalization happens quickly (much less than 1 ms), for a given illumination profile, intensity is a good variable to quantify heating effects.^22^ By adding a reflective and thermally conductive metallic coating to the substrate, such as gold or silver, the excitation intensity can be raised by an order of magnitude to 1000 W/cm^2^ across a 35 µm full width at half maximum (FWHM) gaussian beam.^21^ While an increase in excitation intensity of 20x was found to have a corresponding increase in the localization rate, the reduced quantum yield of photobleaching still keeps the localization rate far below what is achieved at RT. This reduced rate effectively limits the resolution of the technique by limiting how finely the underlying structure can be sampled with precise position measurements within a fixed experiment time, while longer experiment times increase risk of ice contamination and sample damage.

To address this challenge, recent work has sought to find a mechanism to use as an off-switch for FPs at CT. The red FP mApple was found to turn off in response to simultaneous illumination with two colors of light.^23^ Because mApple is not photoactivatable and starts in the bright state, it is not ideal for srCryoCLEM. Fortunately, in this work we identify a similar turn off mechanism for PAmKate, which starts in the dark state. We employ multiple fluorescence techniques to understand the turn off mechanism in PAmKate and better determine how to control the on-off state of molecules to increase localization rate and improve srCryoCLEM fidelity.

## Results

Purified PAmKate was spun coat in a thin film of poly(vinyl alcohol)-(vinylidene alcohol) (PVA-VA) on a glass slide. Most of the molecules were photoactivated by exposure to room lights during the sample preparation. Then, the sample was cooled to near liquid nitrogen temperature in a home-built cryogenic optical stage under a nitrogen atmosphere. Wide-field imaging was done under several illumination conditions, Fig. 1. In one region, the sample was continuously pumped with 594 nm excitation, to pump the singlet-singlet transition as would be done in an imaging experiment. Then, in other regions, the sample was exposed to both 594 and 405 nm light, either one after the other or simultaneously.

**Figure 1:**
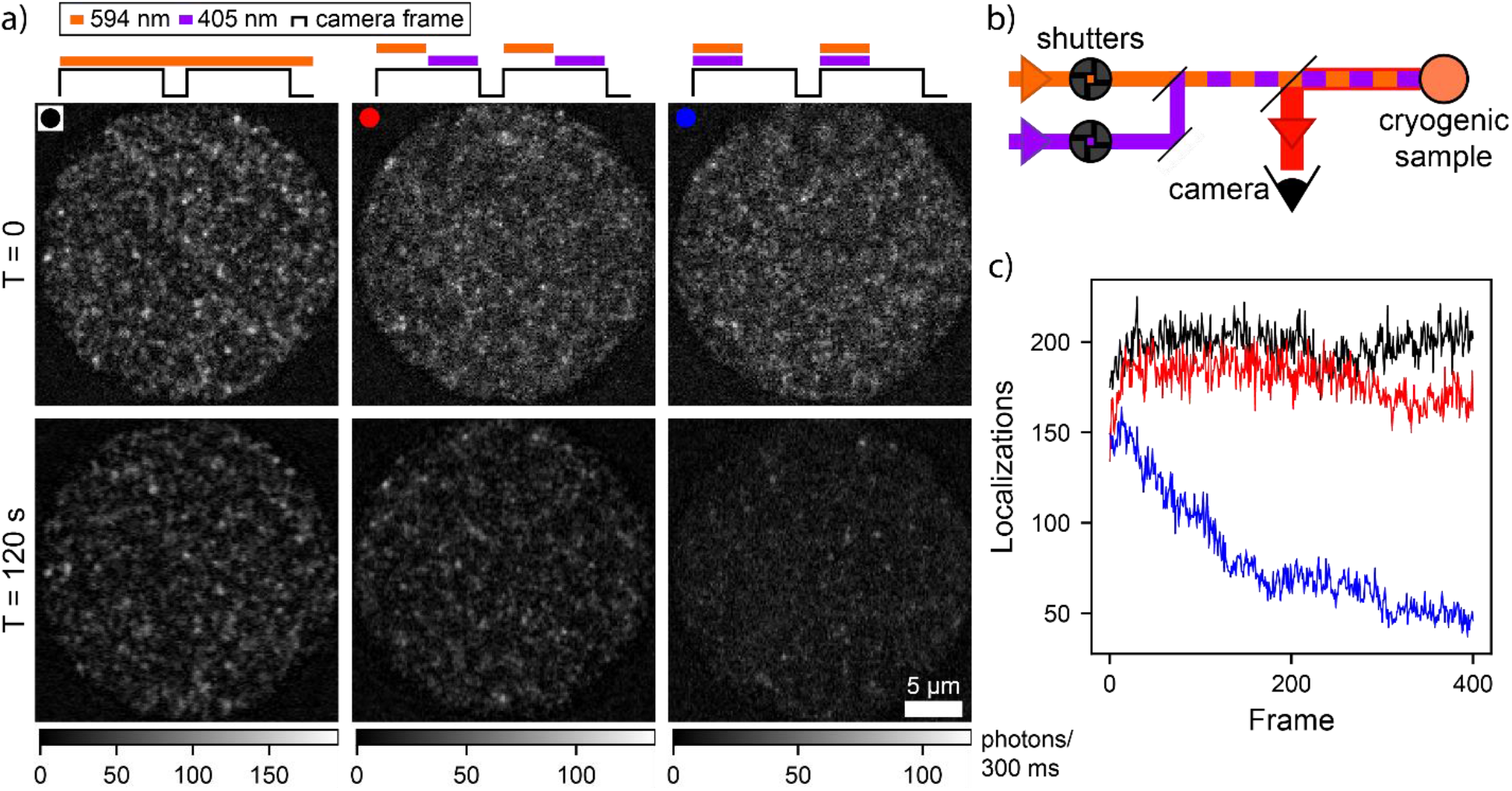
Single-molecule imaging under different illumination conditions at cryogenic temperature. a) First (top) and last (bottom) frames of length 300 ms acquired under different illumination conditions: continuous 594 nm illumination (left), alternating 594 and 405 nm illumination within each frame (middle), and simultaneous 594 and 405 nm illumination for half of each frame (right). b) Experimental configuration. Timing of 405 and 594 nm illumination is controlled with shutters, and fluorescence from the sample is collected on a camera. c) Single-molecule localizations in each frame for continuous 594 nm (black), alternating 594 and 405 nm (red), and simultaneous 594 and 405 nm (blue) illumination. Simultaneous 594 and 405 nm illumination dramatically increases turn-off rate, decreasing the number of emitting molecules in each frame. Average intensity at the sample plane was 300 W/cm^2^ and 80 W/cm^2^ for 594 and 405 nm respectively.

Under continuous 594 nm excitation and alternating 594 and 405 nm, the images are similar. As expected, some photobleaching occurs, but after 120 seconds the field of view is still densely populated with emitters, which would limit the ability to localize new molecules in an imaging experiment. After 120 s of simultaneous 594 and 405 nm illumination, however, the sample looks starkly different. While the density of molecules at T = 0 is similar to the other two conditions, by the end most of the molecules have turned off. Movies of each acquisition are included as Supplementary Video 1.

This result is immediately useful for cryo-PALM imaging with PAmKate, as continuous exposure to 405 and 594 nm can increase localization rate. However, because 405 nm also photoactivates more molecules, the intensity must be controlled to balance turn-on and turn-off to maintain appropriate sparsity. Also, the addition of 405 nm excitation increases background significantly when imaging cellular samples due to autofluorescence. For these reasons, it is valuable to explore the photophysical landscape of PAmKate to search for additional wavelengths or timing schemes which can drive this effect for improved active control.

The difference in turn off rate from alternating *versus* overlapped 405 and 594 nm suggests the existence of a transient state, accessed after excitation by one of the colors, which can be bleached by the other. From this measurement with 300 ms frame times, it can be seen that the lifetime of that state is less than about 100 ms, but shorter time scale information is needed. To explore these questions, we modified our microscope to enable time-resolved measurements of the transient state.

To control the illumination timing faster than a shutter, the digital modulation ports on the 594 and 405 nm lasers were used, with trigger signals generated by dedicated electronics (see methods). The 405 nm laser turns on to a stable power within 5 µs of the trigger, and the 594 nm after 40 µs (see Fig. S1), so we are insensitive to time dependent effects less than about 40 µs. Total fluorescence from a roughly 400 µm^2^ region of high pressure frozen PAmKate (see methods) was collected on an optical receiver, providing much higher time resolution than the camera used in the previous experiment.

To study entry into the transient state, a region of frozen PAmKate was fully photoactivated with 405 nm. Then, the 594 nm excitation was toggled on and off while fluorescence was monitored. The result is shown in Fig. 2a, with quantification in Fig. 2b. With a relatively low 594 nm intensity of 50 W/cm^2^, the fluorescence decreases considerably, by 33% after 5 ms, and is almost fully recovered after sufficient dark time (approximately 10 ms, discussed below in Fig 2c). This means, at steady state, 1/3 of the molecules are shelved in a state which does not emit fluorescence in the collection band at 620 nm under 594 nm excitation. The shelving transient data fits well to a biexponential decay with time constants of 1.4 and 0.35 ms. When the power is increased to 160 W/cm^2^, 50% of the molecules are shelved, and the time constants decrease to 1.0 and 0.23 ms. Interestingly, the addition of 260 W/cm^2^ of broad band near IR radiation (NIR) centered around 750 nm results in a decrease of the fraction shelved (to 45%), a slight increase in the longer time constant (0.98 to 1.03 ms) and repartition to the shorter lifetime (32% to 36%). Notably, the NIR on its own does not produce any fluorescence. This suggests NIR drives molecules out of the transient state, eventually back to the S_0_ ground state where they can be excited by the 594 nm light. This is similar to reverse intersystem crossing (RISC), where NIR is shown to brighten fluorescence by driving molecules out of a dark triplet state back to the singlet ground state,^24^ and suggests this transient state may itself be a triplet state.

**Figure 2:**
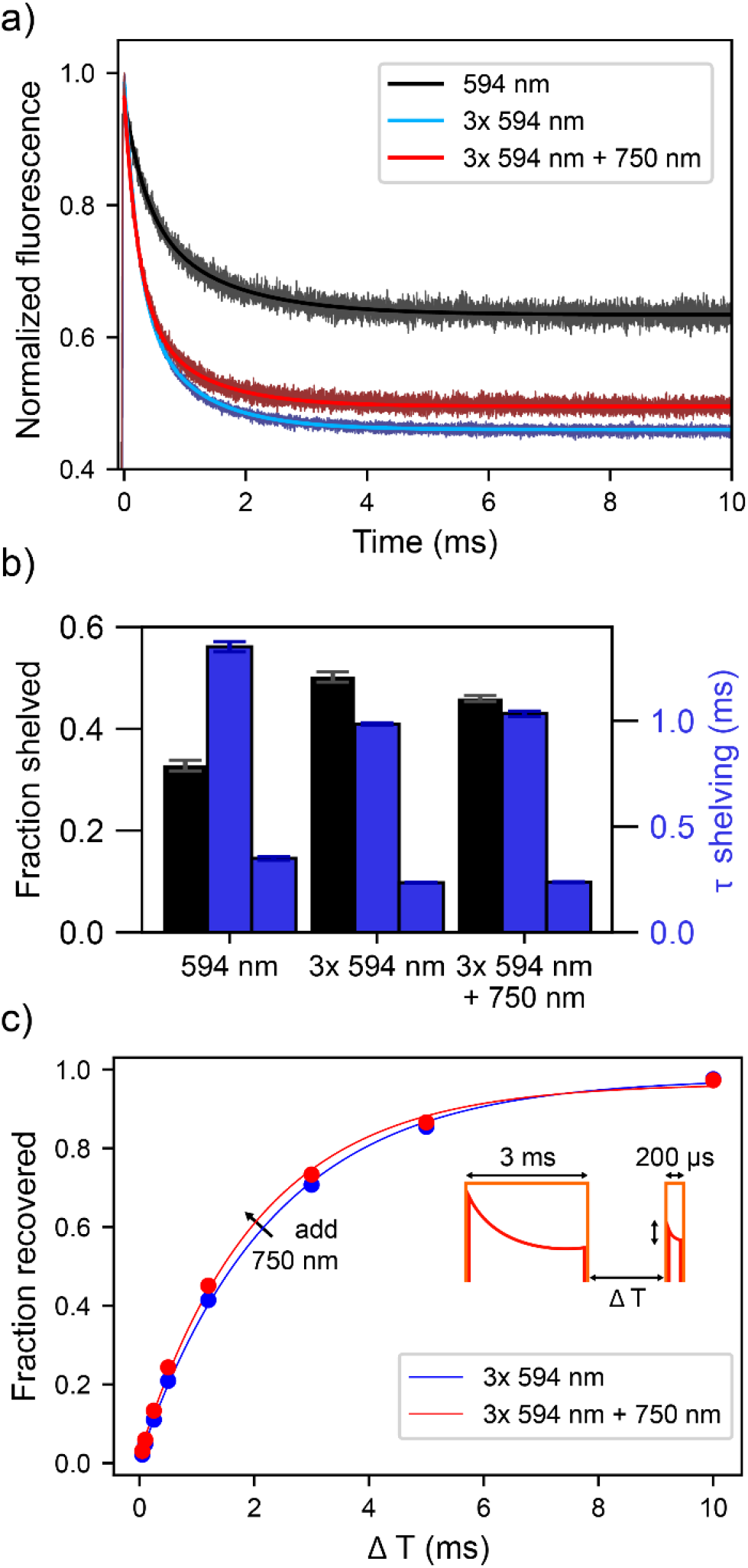
Characterizing population of and recovery from the transient state. a) Shelving into transient state under different illumination conditions, monitored by fluorescence. Curves show 50 W/cm^2^ 594 nm (black), 160 W/cm^2^ 594 nm (blue), and 160 W/cm^2^ 594 nm with 260 W/cm^2^ 750 nm (red) plotted with biexponential fits. b) Fraction shelved (black) and lifetimes from a biexponential fit (blue) for each condition, with error bars showing standard deviation of fit. c) Fraction of molecules recovered from transient state as a function of time for 160 W/cm^2^ 594 nm (blue) and 160 W/cm^2^ 594 nm with 260 W/cm^2^ 750 nm (red). Data are shown as points, with standard deviation smaller than each point, and single exponential fits are solid lines. Inset shows timing of excitation (orange) and fluorescence (red).

By following an initial 3 ms long shelving pulse with a 200 µs probing pulse with a variable time delay, ΔT, the rate of recovery from the transient state can be measured. The fraction recovered is estimated by measuring how much fluorescence, lost in the first pulse, has recovered by the second probing pulse. This is shown, along with single exponential fits in Fig. 2c. With only 594 nm illumination, the recovery proceeds with a lifetime of 2.3 ms, regardless of intensity (see Fig. S2). However, with the addition of 260 W/cm^2^ at 750 nm (on continuously during shelving and recovery), the recovery time shortens slightly to 2.1 ms, consistent with NIR driving molecules out of the transient state.

If this state is indeed the bleachable transient which leads to turnoff of the emission, it would be expected that bleaching with 405 nm should occur on a similar timescale. To investigate this, a new experimental scheme was utilized which features pairs of pulses and allows for the control of the time between 594 nm and 405 nm interactions. As shown in Fig. 3a, each 200 µs pulse of 594 nm is followed by a 200 µs pulse of 405 nm with a variable delay time between them, ΔT. Nine pulse pairs occur during one camera frame. Every four seconds, the delay (ΔT) is changed to another value in a random pattern. Fluorescence is collected on a camera, and for each four second window, the change in fluorescence brightness (ΔF) is measured. This change is divided by the average of that window (F) to yield a fractional change in population. Fig. 3b shows a small portion of raw data, showing steeper slopes for shorter delays. Results for this experiment averaged over many repetitions and several regions of sample are shown as red dots in Fig. 3c, with error bars representing the standard error of the mean. ΔF/F starts negative, and its magnitude decreases with increasing ΔT. When the 405 nm pulse arrives shortly after the 594 nm pulse, the sample bleaches rapidly. As the delay increases, the bleaching effect goes away, even turning slightly positive for large ΔT. A single exponential fit provides a decay time of 1.8 ms, similar to the 2.3 ms recovery time of the transient state measured in the experiment above (Figure 2c). This suggests that the transient state observed in the fluorescence recovery experiment is the same state which can be bleached by 405 nm, leading to the enhanced bleaching rate under simultaneous illumination. In future work, this time scale could be leveraged to provide further control over the on-fraction of PAmKate. If the 405 and 594 nm excitations are separated by more than a few ms, photoactivation will dominate and the density of molecules will increase. If the delay is short, photobleaching will be enhanced, and the density of molecules can be decreased. Thus, the timing of the 405 and 594 nm excitation can dictate if more PAmKate molecules are being turned “on” or “off.” As a control, the 405 nm pulse was replaced with another 200 µs pulse of 594 nm. ΔF/F was divided by two to account for the additional 594 nm power. In this case, no change in ΔF/F is observed as ΔT varies, showing 594 nm does not bleach the transient state, Fig. 3c.

**Figure 3:**
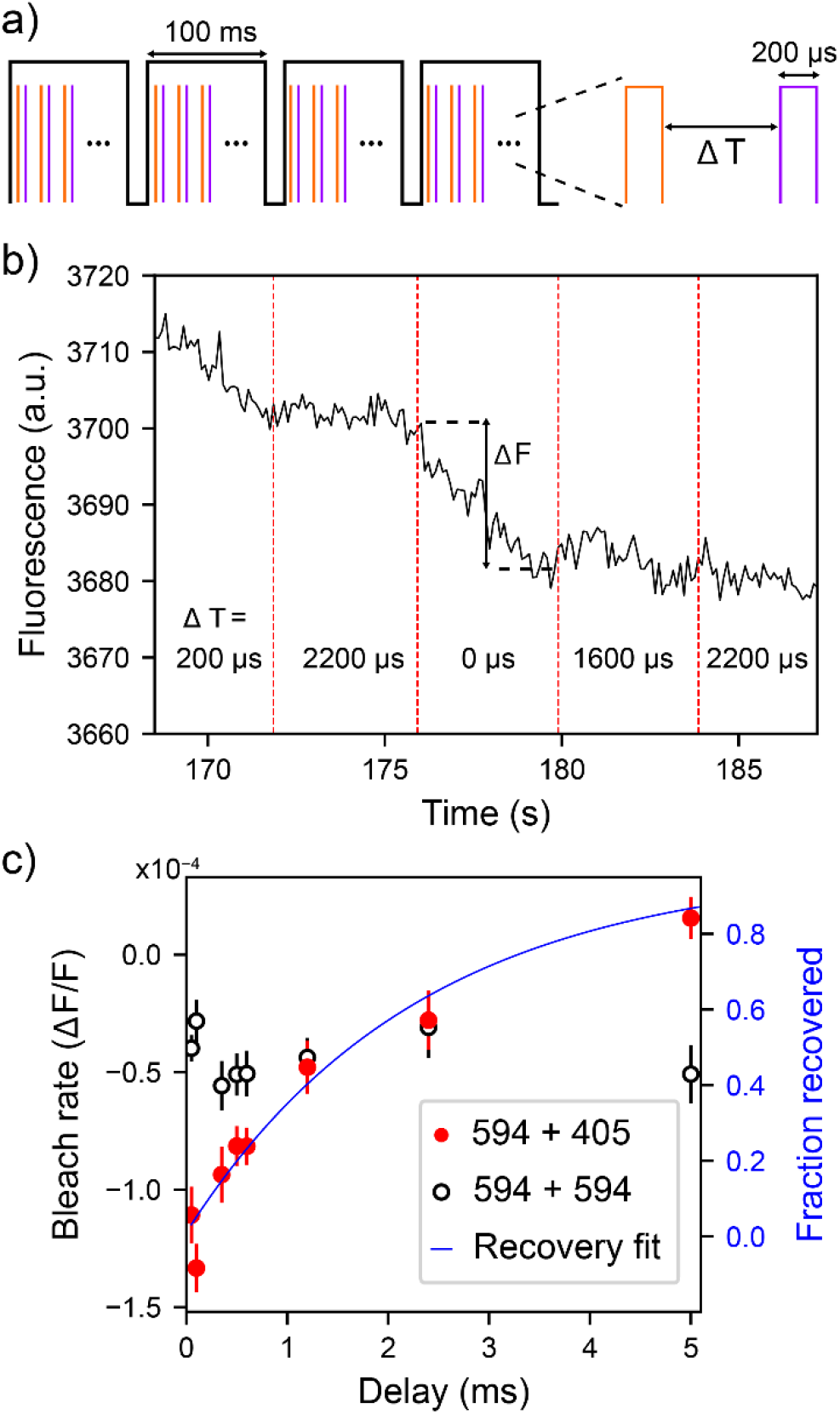
Characterizing bleaching from the transient state. a) Experimental timing diagram. 200 µs pulses of 594 nm (orange) are followed after delay ΔT by 200 µs pulses of 405 nm (blue). During each laser on time, intensity at the sample of 594 and 405 nm was 500 and 200 W/cm^2^ respectively. This repeats 9 times for each 100 ms camera frame (black). b) Raw fluorescence data averaged over the illumination spot. The slope of bleaching changes as the delay between pulses is changed. c) Bleach rate versus delay time for 405 nm (red circles) and 594 nm (black circles) second pulse. 594 + 594 nm is scaled by a factor of ½ as it has double the 594 nm flux. Error bars represent standard error of the mean. The exponential fit to the 594 nm-only recovery from a separate experiment shown in Fig. 2c is plotted as a blue line. The fit to the bleach rate data is presented in Fig. S3 for comparison.

With the lifetime of the transient state identified, we next turn to characterization of the response to different wavelengths, i.e., we seek to establish an action spectrum of bleach from the transient state. While the 405 and 594 nm lasers can be quickly modulated by a digital signal, the other light sources cannot. We therefore measure the bleaching enhancement for a variety of wavelengths with an experimental setup like the one used for the single-molecule imaging diagrammed in Fig. 1b, in this case averaging fluorescence over a region of high pressure frozen PAmKate instead of imaging single molecules. Shutters are used to either overlap or alternate exposure to 594 nm and a second color (λ_2_) within each frame, toggling between the conditions every few seconds. The bleaching rate between these conditions is then compared, with results shown in Fig. 4a.

**Figure 4:**
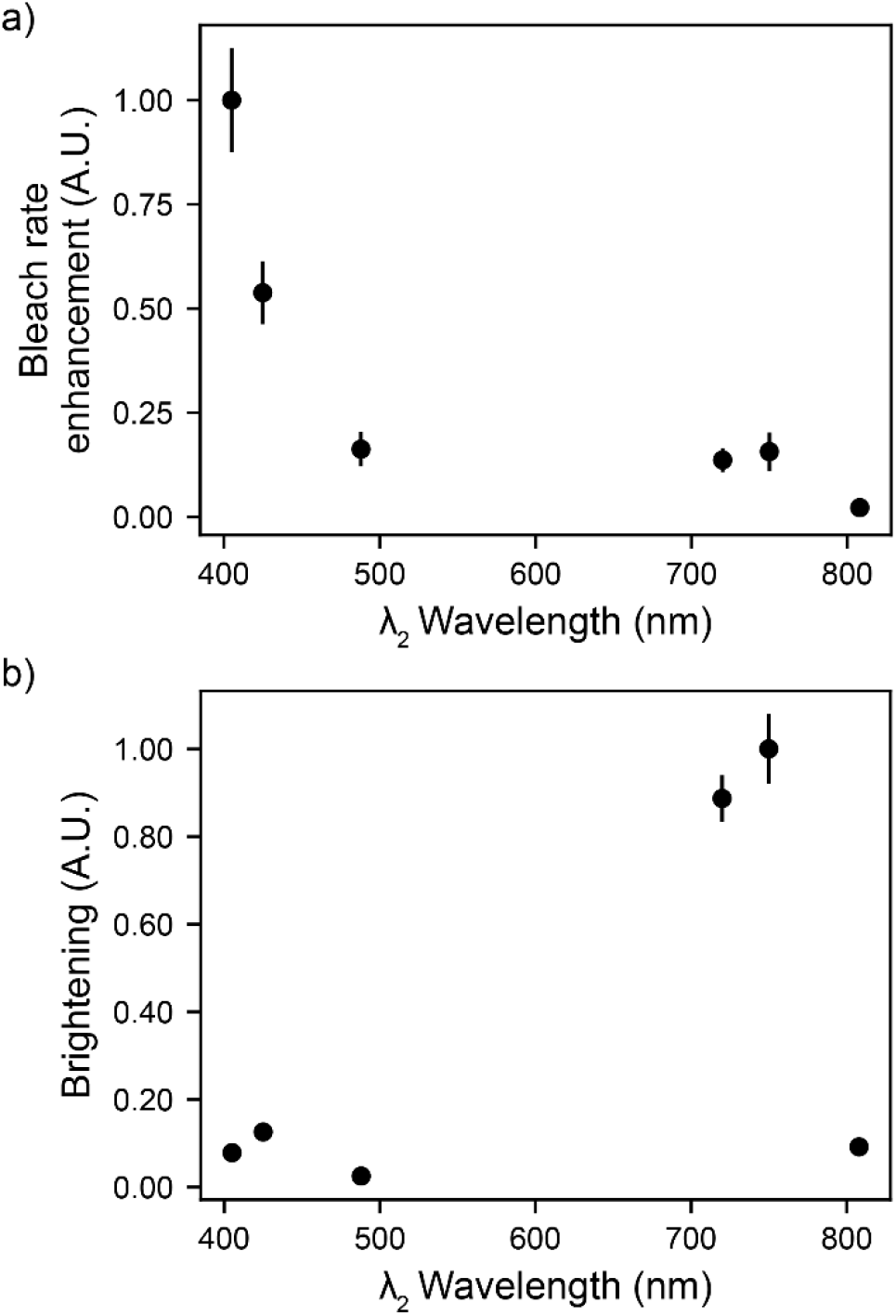
Action spectra of the transient state. a) Bleach rate enhancement is defined as the slope of bleaching with 594 nm and λ_2_ overlapped minus the slope while alternated, normalized to λ_2_ = 405 nm. b) Brightening is defined as the change in fluorescence from the last point of the alternating condition to the first point of the overlap condition, normalized to λ_2_ = 750 nm. Error bars are standard deviation. λ_2_ powers were set for equal photon flux at the sample plane where possible. For 750 and 808 nm, the values are scaled to account for the reduced photon flux. Raw data is shown in Fig. S4.

Of all the wavelengths tested, 405 nm showed the largest response, followed by 425 and 488 nm. Since 561 and 594 nm excitation drives molecules into the transient state, their ability to bleach from the transient state could not be quantified in this experiment. However, we know from the flat response of bleach rate vs delay for two pulses of 594 nm (Fig. 3c black circles) there is not a large effect at this wavelength. Interestingly, another small peak is observed in the NIR with significance beyond the error bars, with 720 and 750 nm exhibiting 15% of the bleach rate enhancement of 405 nm. Both the NIR and 405 nm, when interacting with the transient state populated after exposure to 594 nm, increase fluorescence by driving molecules back to the ground state (Fig. 4a) and increase the rate of bleaching. However, the relative magnitudes of these effects are different, with NIR driving relatively more brightening than bleaching compared with 405 nm. This suggests that while these wavelengths may be interacting with the same state, they are driving molecules to different excited states with different probabilities to return to the active state or bleach.

In the process of these measurements, we observed a secondary effect that is important to consider for experimental design and imaging applications with PAmKate. Similar to other FPs, the photobleaching curve does not fit well to a single exponential, regardless of the wavelengths involved. Specifically, there appear to be two primary effects with a short and long timescale, as occurs with many other photobleaching effects with fluorescent proteins.^25^ While photobleaching, including bleaching from the transient state, dominates at longer times, early times are dominated by driving molecules into a long-lived dark state. This is reminiscent of photoswitching observed in GFP variants at CT, attributed to photoinduced protonation of the chromophore.^26^ Recent work has shown that rsEGFP2 photoswitching proceeds through a different mechanism at CT than at RT, resulting in multiple dark states without any large conformational changes.^17^ It is possible that the photoswitching observed here in PAmKate proceeds through a similar mechanism. These photoswitched molecules can be returned to the active state by exposure to 405, 425, and 488 nm, but not by the NIR (Fig. S5). Because the initial photoactivation of PAmKate is thought to proceed through decarboxylation,^27^ it is unlikely this photoswitching represents a return to the precursor molecule; instead, it points to an additional long-lived dark state.

For the purposes of this study on the bleachable transient state, it is important to consider the establishment of equilibrium between the active and dark states. For all the experiments presented in this paper, data was only considered for analysis after sufficient time had passed to stabilize this equilibrium. For the purposes of imaging, it is important to note that 405 nm intended for photoactivation of new PAmKate molecules will also switch on some molecules which had already been imaged, which will result in overcounting if not properly analyzed.

To demonstrate how the active control mechanism described in this paper can be used to improve cryogenic super-resolution, we applied this method to imaging a particular protein fusion in *Caulobacter crescentus*. This freshwater bacterium is interesting to developmental biologists due to its asymmetric division. Here, we use a strain which contains an overexpression of crescentin (CreS) fused to PAmKate (see methods). CreS is necessary for *C. crescentus* to adopt its characteristic curved structure; without it the cells grow as straight rods.^28^ CreS has been found to form structured intermediate filaments along the inner edge of the cell, and it is thought this structure applies its geometry to the cell.^29-31^

*C. crescentus* cells expressing PAmKate-CreS fusion constructs were frozen on custom silver grids^21^ and imaged with a mixture of 405 and 561 nm excitation (see methods). Importantly, cells were not preactivated with 405 nm excitation and the majority of PAmKate molecules started in a dark state. Figure 5 shows both a diffraction limited overlay as well as a representation of the single-molecule localizations achieved after just 12 minutes of imaging. Previous work by the authors, before the knowledge of reduced heating on metallic grids and the exploitable PAmKate photophysics, achieved lower density of localizations after 3 hours of imaging.^11^ The results shown here and in Fig. S7 demonstrate that the intermediate state identified in PAmKate can be exploited for improved control under cryogenic conditions, yielding higher localization rates and improved super-resolution reconstructions.

**Figure 5:**
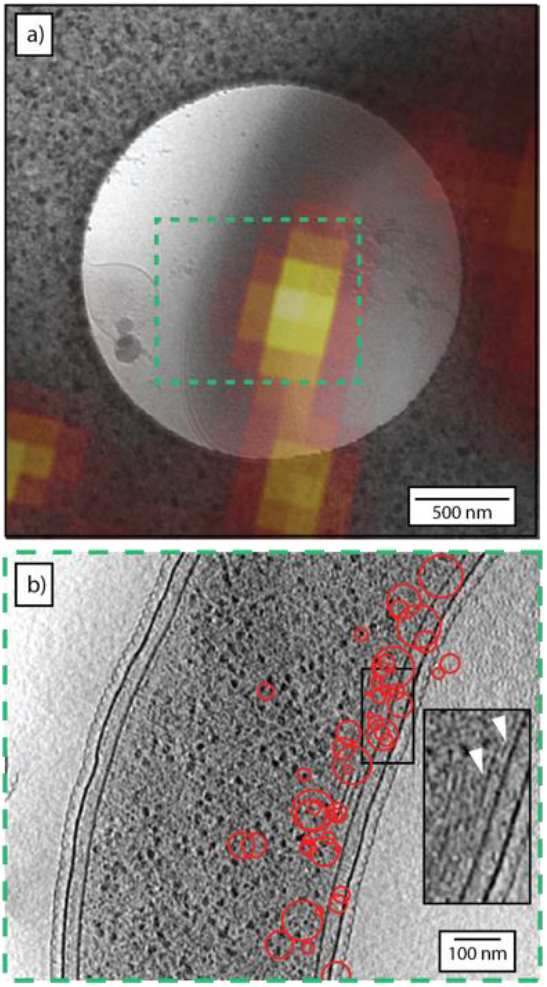
Example of srCryoCLEM of PAmKate-CreS. a) Diffraction-limited overlay of fluorescence (heat map) and low-magnification cryogenic electron micrograph (gray) of *C. crescentus* cell expressing PAmKate-CreS fusion construct. b) Overlay of single-molecule fluorescence localizations (red circles) and a central slice from a tomographic reconstruction of the region bounded in a) by the dashed green box. The radius of each circle is the experimentally determined localization precision. Inset shows fibrils along the ventral side of the cell similar to those that have been attributed to CreS previously.^30^

## Discussion

Combining the experimental results presented above with results from previous work, we propose a model of the photophysical landscape of PAmKate at CT, shown in Fig. 6. This model is consistent with our observations but is not the only possible interpretation. When there is ambiguity about whether two processes lead to one or multiple distinct states, for simplicity we draw it as one state unless our observations show definitively otherwise.

**Figure 6:**
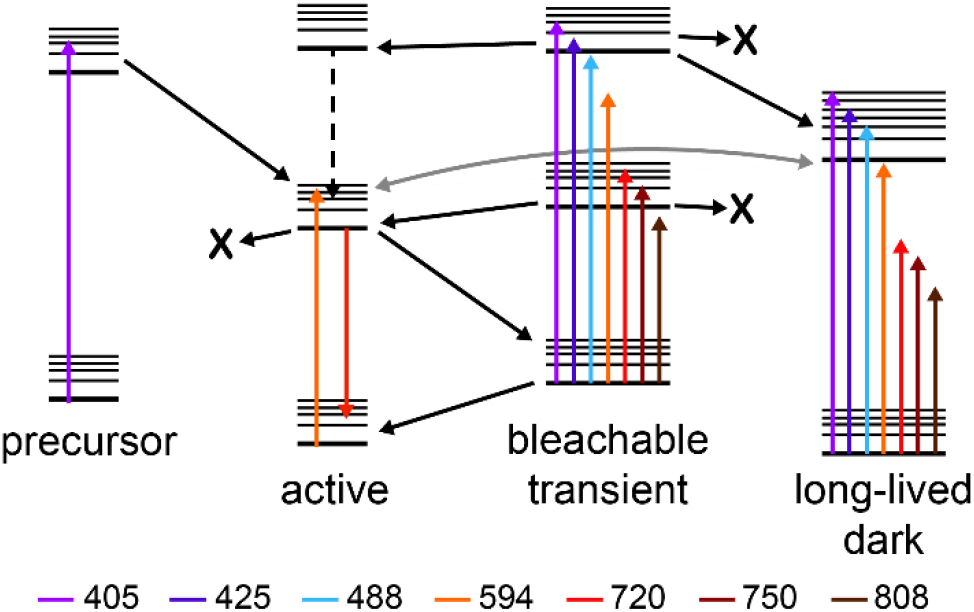
Diagram of observed photophysical states at 77K. Each arrow represents a transition inferred from an experiment in this paper or a previous work, colored according to wavelength. Black and gray arrows have unknown wavelength or no emission. X represents a molecule which has irreversibly photobleached.

The model contains four distinct ground states, each with their own electronic structure. First, the precursor state is dark in the collection channel of 600-700 nm. 405 nm excitation can convert precursor to active, even at 77 K. This key feature makes PAmKate a good candidate for cryo-PALM, but without a turn-off mechanism, the prolonged on times of active molecules limit throughput and sampling. The active state, once excited by 594 nm, has some probability to emit fluorescence which can be used for localization. It also can irreversibly photobleach (with a lower probability than at RT) or photoswitch into a long-lived dark state (gray arrow). Molecules in this dark state can be photoswitched back to the active state with blue/UV excitation (gray arrow), allowing for continued localization improving precision, but potentially leading to overcounting. Finally, excited active state molecules may enter a transient state with a lifetime of ∼2 ms. The biphasic nature of this shelving process suggests multiple species are formed, either independently or sequentially.

Molecules in the transient state are susceptible to blue/UV and NIR irradiation. When excited by 405 nm, molecules have some probability to return to the active state, shortening the lifetime of the transient and allowing for excitation and collection of more fluorescence. They also have some probability to irreversibly photobleach, providing the desired turn-off mechanism. Finally, they may also enter a long-lived dark state from the excited transient state (Fig. S8 arrow 10). When illumination is changed from overlap to alternating, the upward slope seen in Fig. S4d is due to a change in equilibrium between this dark state and the active state (Fig. S8 arrow 11). This dark state is drawn in Fig. 6 as the same long-lived dark state described above for simplicity, as described at the beginning of the section.

Under 720 and 750 nm illumination, molecules in the transient state have some probability to return to the active state or permanently bleach. Because the ratio of brightening to bleaching is different for molecules exposed to NIR compared to blue/UV, they must be driving molecules to different excited states.

While the identity of this transient state cannot be determined from the experiments here, comparison with literature suggests a few options. Commonly invoked identities for transient states in FPs include triplets, protonation, redox change, or conformational changes.^32, 33^ As described previously, conformational changes are unlikely in a cryogenic environment. A triplet state is a possible candidate, supported by the observation of what would be called reverse intersystem crossing.^34, 35^ Recent work on GFP mutants found a triplet lifetime of 5 ms at RT and 20 ms at CT, with an absorption peak around 900 nm.^36, 37^ RT transient absorption spectroscopy of three red FPs identified a transient state with a lifetime of µs to ms and an absorption peak of 730 nm. The authors assigned this state to a dianionic-radical chromophore.^38^ Based on these comparisons, we believe the transient state observed here may be a triplet or dianionic radical, likely formed from the triplet. To unequivocally assign the identity of this state, further experiments are necessary. Following the blueprint of Refs. ^36, 37^, transient absorption spectroscopy of PAmKate, first at room temperature and then at cryogenic temperature, would provide useful insights. Room temperature measurements facilitate introduction of chemicals to promote or suppress certain species (such as triplet quenchers) which can aid in identification, and comparison to cryogenic temperature connects these observations to the states populated in cryogenic imaging experiments.

Recent work at room temperature found simultaneous illumination with near-infrared and the excitation wavelength reduced photobleaching of GFP variants, but increased photobleaching of red fluorescent proteins in certain environmental conditions.^39^ Since the environmental conditions in their experiment were not well controlled, the authors do not propose a mechanism, but it is possibly related to the effect observed in this work.

Here, no shelving to the transient state was observed for PAmKate samples at RT (Fig. S6). This means either the lifetime of the state is much shorter or the quantum yield of shelving is much lower at RT compared to CT, resulting in a response below the detection limit. To identify high performing cryogenic fluorophores, it is necessary to look at their behavior at CT. The different temperature and environment mean active control pathways accessible at CT may not be accessible at RT and vice versa.

While the primary focus of this work is on photophysical control of active PAmKate molecules at CT, it is worth noting that similar multistep pathways may also exist in the precursor state, as observed at room temperature in the green fluorescent protein mEos4b.^40^ While at present long on times limit sampling, it will eventually be important to consider the photophysics of the precursor to maximize labeling efficiency and the resolution of the recovered structure.

## Conclusion

While much work has been directed towards optimizing fluorescent labels and imaging conditions for RT super-resolution microscopy, the field of srCryoCLEM is only beginning to explore the possibilities. Identifying photoactivatable and photoswitchable labels is a necessary start, but the challenges and opportunities of imaging at CT are broader. The low temperature condition means certain states have longer lifetimes, and multi-step pathways which require prohibitively high photon fluxes at RT may be accessible. By leveraging these pathways and taking advantage of higher emitted photon counts and the minimally perturbative fixation method of plunge freezing, cryogenic super resolution should eventually exceed what is possible at RT, all while remaining compatible with cryo electron tomography.

While the focus of this work is on leveraging PAmKate photophysics for cryo-PALM and single-molecule imaging techniques, it is also worth noting the high population, saturability, and lifetime of the transient state make it a good candidate for use in nonlinear structured illumination microscopy.^41^ Regarding cryo-PALM, the results of this work can be immediately used to improve active control in srCryoCLEM with PAmKate, as demonstrated in Fig. 5. By adding a continuous low fluence of 405 nm or a high fluence of 750 nm, molecules can be brightened and blinking dynamics increased. When imaging samples on bio-compatible gold EM grids, a higher intensity of 750 nm can be used for the same heating as 405 nm, for roughly equal rates bleaching from the transient. Further, by controlling the delay between pulses of 594 and 405 nm, the ratio of photoactivation to photobleach can be tuned.

The transient state identified in this work, which provides access to bleaching and brightening pathways, is likely to exist for other related FPs. Further work on various mutants could inform which residues are key to promoting or stabilizing this state, leading to opportunities to engineer FPs with desirable traits for specific imaging challenges. This work provides an experimental blueprint to search for multicolor stepwise control effects in FPs at cryogenic temperatures.

## Methods

### Protein expression and purification

Purification was performed according to Cranfill *et al*. with minor modifications.^42^ Briefly, One Shot TOP10 Chemically Competent *E. coli* (Thermo Fisher, Cat# C404003) were transformed with pBAD/HisB-PAmKate (Plasmid #32691, Addgene)^43^ and grown to OD600 0.6 in LB medium containing 0.1 mg ml−1 ampicillin at 37 °C. Heterologous protein expression was then initiated with 0.02% w/v L-Arabinose (Sigma A3256) overnight at 30 °C. Cells were collected by centrifugation (4000x g, 10min) and frozen. For purification, cells were resuspended and lysed in BugBuster® Master Mix Protein Extraction Reagent (Millipore Sigma, Cat# 71456) at room temperature following the manufacturers recommendation. Lysate was cleared by centrifugation (15,000x g, 10min) and the soluble fraction of His-tagged proteins was purified using His SpinTrap columns (Cytiva Cat# 28401353) following the manufacturer’s protocol. The eluate protein from NiNTA columns was buffer exchanged into PBS pH 7.4 and concentrated with a 3 kDa Amicon® Ultra-4 Centrifugal Filter Units (Millipore Sigma Cat# UFC800308). Proteins were frozen and stored at -80 °C.

### Single-molecule imaging

Purified PAmKate was diluted in PBS and 0.5 % PVA-VA and spin coated onto a glow-discharged glass coverslip. Imaging was done in a home-built open cup cryostat. The sample was placed on a copper pedestal mounted in an insulated stainless-steel cup. Liquid nitrogen was added to the cup, cooling the pedestal and the sample to near 77K. The sample was imaged with a Nikon CFI Plan Achromat LWD IMSI 100XC (NA 0.85) imaging through a 1 mm glass window. 594 nm (Coherent OBIS 594-100 LS) and 405 nm (Coherent OBIS 405-100 LX) lasers were coupled into a 200 µm square-core fiber (Newport De-Speckler SQR200) which was cropped by an iris imaged onto the sample to provide flat illumination over a field of view of approximately 25 µm diameter. Average intensity at the sample plane was 300 W/cm^2^ and 80 W/cm^2^ for 594 and 405 nm respectively. Fluorescence from the sample was filtered by a dichroic beamsplitter (Chroma 59003bs), a 594 notch filter, a 594 long pass filter, and a 610/60 band pass filter before detection on a camera (Andor iXon Ultra 897). For the overlap condition, the fire signal from the camera (300 ms exposure) was sent to an Arduino Uno, which sent a pulse to open shutters for both lasers for 130 ms. For the alternating condition, the pulse to the 405 nm shutter was delayed 150 ms. Hardware was controlled using PYMEAcquire.^44^ Drift correction was done by cross correlation, and background subtraction by 12 µm top hat filtering. To estimate the number of emitters on in each frame, localization was done using the LatGaussFitFR module in PYME.^44^

### Sample preparation and data analysis for srCryoCLEM demonstration

Cells with a creS-PAmKate plasmid were induced overnight, deposited on custom silver-coated finder grids (Ref. ^21^) and plunge frozen. The grids were loaded on a cryogenic fluorescence microscope similar to that previously described^23^ and illuminated with gaussian profiles with peak intensity of 770 and 30 W/cm^2^ of 561 and 405 nm respectively. At the particular cell in Figure 5 and S7, selected for correlative analysis because it is mostly over a hole in the EM grid, the illumination intensity was 600 and 20 W/cm^2^ of 561 and 405 nm respectively. Blinking data was collected with this intensity for about 7 minutes, after which the 405 nm intensity was doubled for an additional 5 minutes, as blinking events were viewed to decrease in frequency.

After fluorescence imaging, the grids were transferred to and imaged in a 300kV electron microscope (Titan Krios ThermoFisher) with a K3 Gatan direct electron detector and energy filter. Cryogenic tilt-series were collected with a dose-symmetric scheme varied between -60° and 60° and a 2° step size. A spot-size of 5 and a target defocus of 10 μm were applied. The pixel size was 4.625 Å and the magnification was 19500x. The total electron dose was limited to ∼120 e/Å^2^ over the acquired tilt-series. Alignment of the tilt-series was achieved using selection of (non-gold) fiducials present on the grids located peripherally to *Caulobacter* cells (small ice particles, gird hole defects) and the Etomo package.^45^ Additional low-magnification (pixel size 36.98 Å) micrographs were taken after the acquisition of tomography data. These low-magnification micrographs are used to register optical and electron microscopy datasets.

Single molecules were localized following the protocol described in Refs ^11, 21^. Briefly, single molecules were merged using a combination of Matlab and Thunderstorm. Matlab was used to manually identify single emitters from step-wise dynamics observable in the fluorescence intensity and estimate the background often from a combination of autofluorescence and other neighboring emitters. This background was subtracted and the resulting fluorescence PSF was fit in each frame using Thunderstorm and then merged into a single localization with the uncertainty in position estimated from the experimental standard error on the mean of the frame localizations. The single-molecule localizations were then registered to the CryoET data using a series of affine transformations. First the optical images were registered to a low magnification EM image where ∼12-16 holes in the holey support film are visible. The centers of these holes, visible in both imaging modalities, were used to compute a projective transformation carrying the optical data into the low-magnification EM space. This low-magnification EM space was then registered with the CryoET space by identifying unique features, such as gold beads and ice contamination, visible in both the low-magnification image and the z-projection of the CryoET reconstruction. From these control point pairs a similar transformation was computed to carry the low-magnification EM micrograph to the CryoET space. Application of the projective and similar transformations sequentially to the optical data carries the single-molecule fluorescence localizations into the CryoET space.

### HPF sample preparation

Purified PAmKate was diluted (1:5) into a solution of PBS (20 mM, pH 7.4) and 30% (v/v) ethylene glycol. An aliquot of 10 µL was added to the 0.2 mm side of a glow discharged Type A 6 mm gold planchette and sandwiched with the flat side of a Type B 6 mm gold planchette. The sandwich planchette was immediately transferred to a high-pressure freezing system (Leica EM ICE, Leica Microsystems) to generate a thick layer (∼0.2 mm) of frozen protein. After cryogenic freezing, the gold planchettes were stored in liquid nitrogen.

### Shelving to transient state

A planchette of high pressure frozen PAmKate was loaded onto the setup described above, with minor changes. A supercontinuum laser (Leukos Rock 400) with a tunable filter (Leukos Bebop) was used to provide light centered around 750 nm with a bandwidth of about 80 nm and an average intensity at the sample of 260 W/cm^2^. 594 nm intensity was set to 55 or 160 W/cm^2^. The De-Speckler was not used, and illumination was cropped to approximately 10 µm with an iris. A different objective (Nikon M Plan 100/NA 0.80 ELWD) was used, and a 90/10 beamsplitter was introduced after the filters, sending 90% of the light onto an optical receiver (Newport 2001). To quickly toggle the 594 nm laser, a function generator (Hewlett Packard 33120A) was used to produce a 25 Hz square wave which was passed to the digital modulation port of the laser controller, repeatedly turning the laser on for 20 ms and off for 20 ms. Fluorescence signal collected on the receiver was read out with a digital oscilloscope (Picoscope 4262) synchronized to the function generator. 2000 to 4000 repetitions are averaged to produce the plots in Fig. 2a.

### Recovery from transient state

The experimental configuration was as described for the shelving experiment with a few changes. The function generator was run at 40 Hz, triggering a delay generator (Stanford Research Systems DG535) to produce a 3000 µs pulse with zero delay and a 200 µs pulse with variable delay between 50 µs to 10 ms. Both pulses were sent to the 594 nm laser. The oscilloscope averaged 100 traces at a given delay, after which a new delay was programmed.

### Time resolved bleaching from transient state

A planchette of high pressure frozen PAmKate was loaded onto the setup described above in the section on single molecule imaging with some modifications. The fire signal from the camera was used to trigger 9 cycles of 90 Hz square wave from the function generator synchronized with the camera frame rate. The output was used to trigger the delay generator to produce a 200 µs pulse, sent to the 594 nm laser, and another 200 µs pulse sent to the either the 594 or 405 nm laser after a programmed delay. During each laser’s on time, intensity at the sample of 594 and 405 nm was 500 and 200 W/cm^2^ respectively. The delay was updated every 6 seconds. Fluorescence collected on the camera was averaged over the illumination spot, and the mean (F) and slope (ΔF) of the signal are calculated for each condition.

### Action spectrum

A planchette of high pressure frozen PAmKate was loaded onto the setup described above in the section on single molecule imaging, with the addition of 425 nm (Omicron LuxX 425-120), 488 nm (Coherent Sapphire 488-100 CW), and 808 nm (Philips PLA 1520-808) lasers. The tunable filter is used on the output of the supercontinuum source to provide light centered around 720 and 750 nm. When possible, the output of each laser was set to provide 2*10^20^ photons/s/cm^2^ at the sample. At 750 and 808 nm sufficient power was not available, so their values on the plot were scaled accordingly. The fire signal from the camera (300 ms exposure) was sent to an Arduino Uno which sent a pulse to a shutter for the 594 nm laser and a shutter for another laser. For the overlap condition, both shutters open for the first 130 ms of the frame, and for the alternating condition the 594 nm shutter opens first, and the other shutter opens 150 ms later. The condition was changed from overlap to alternating every 6 seconds, and the bleaching slopes were compared between the two conditions for a given wavelength. Then, the second shutter was moved, and the experiment repeated for a different wavelength.

## Acknowledgments

This work was supported in part by the National Institute of General Medical Sciences grant no. R35GM118067 (W.E.M.). P.D.D. was supported in part by the Panofsky Fellowship at the SLAC National Accelerator Laboratory as part of the Department of Energy Laboratory Directed Research and Development program under contract DE-AC02-76SF00515, and by grant 2021-234593 from the Chan Zuckerberg Initiative DAF, an advised fund of Silicon Valley Community Foundation. T.B.A is supported by Schmidt Science Fellows, in partnership with the Rhodes Trust and was supported by Wellcome (102164/Z/13/Z).

## Associated content

**Supplementary Figures**. Additional figures characterizing the experimental system. Additional data discussed in the text. Annotated state diagram with a table of supporting evidence.

**Supplementary Video**. Videos of PAmKate single molecules under each condition described in the text.

## References

(1) Albert, S.; Wietrzynski, W.; Lee, C. W.; Schaffer, M.; Beck, F.; Schuller, J. M.; Salome, P. A.; Plitzko, J. M.; Baumeister, W.; Engel, B. D. Direct visualization of degradation microcompartments at the ER membrane. Proceedings of the National Academy of Sciences of the United States of America 2020, 117 (2), 1069–1080. DOI: 10.1073/pnas.1905641117 [doi] (acccessed Jan 14).

(2) Dai, W.; Chen, M. Y.; Myers, C.; Ludtke, S. J.; Pettitt, B. M.; King, J. A.; Schmid, M. F.; Chiu, W. Visualizing Individual RuBisCO and Its Assembly into Carboxysomes in Marine Cyanobacteria by Cryo-Electron Tomography. Journal of Molecular Biology 2018, 430 (21), 4156–4167. DOI: 10.1016/j.jmb.2018.08.013.

(3) Wietrzynski, W.; Schaffer, M.; Tegunov, D.; Albert, S.; Kanazawa, A.; Plitzko, J. M.; Baumeister, W.; Engel, B. D. Charting the native architecture of Chlamydomonas thylakoid membranes with single-molecule precision. Elife 2020, 9, e53740.

(4) Russo, C. J.; Dickerson, Joshua L.; Naydenova, K. Cryomicroscopy in situ: what is the smallest molecule that can be directly identified without labels in a cell? Faraday Discussions 2022, 240 (0), 277-302, 10.1039/D2FD00076H. DOI: 10.1039/D2FD00076H.

(5) Wang, Q.; Mercogliano, C. P.; Löwe, J. A Ferritin-Based Label for Cellular Electron Cryotomography. Structure 2011, 19 (2), 147–154. DOI: 10.1016/j.str.2010.12.002 (acccessed 02/09).

(6) Fung, H. K. H.; Hayashi, Y.; Salo, V. T.; Babenko, A.; Zagoriy, I.; Brunner, A.; Ellenberg, J.; Müller, C. W.; Cuylen-Haering, S.; Mahamid, J. Genetically encoded multimeric tags for subcellular protein localization in cryo-EM. Nature Methods 2023, 20 (12), 1900–1908. DOI: 10.1038/s41592-023-02053-0.

(7) Hampton, C. M.; Strauss, J. D.; Ke, Z.; Dillard, R. S.; Hammonds, J. E.; Alonas, E.; Desai, T. M.; Marin, M.; Storms, R. E.; Leon, F.; et al. Correlated fluorescence microscopy and cryo-electron tomography of virus-infected or transfected mammalian cells. Nature Protocols 2017, 12 (1), 150–167. DOI: 10.1038/nprot.2016.168.

(8) Schwartz, C. L.; Sarbash, V. I.; Ataullakhanov, F. I.; McIntosh, J. R.; Nicastro, D. Cryofluorescence microscopy facilitates correlations between light and cryo-electron microscopy and reduces the rate of photobleaching. Journal of microscopy 2007, 227 (2), 98–109.

(9) Klein, S.; Wimmer, B. H.; Winter, S. L.; Kolovou, A.; Laketa, V.; Chlanda, P. Post-correlation on-lamella cryo-CLEM reveals the membrane architecture of lamellar bodies. Communications Biology 2021, 4 (1), 137. DOI: 10.1038/s42003-020-01567-z.

(10) Chang, Y. W.; Chen, S.; Tocheva, E. I.; Treuner-Lange, A.; Lobach, S.; Sogaard-Andersen, L.; Jensen, G. J. Correlated cryogenic photoactivated localization microscopy and cryo-electron tomography. Nature methods 2014, 11 (7), 737–739. DOI: 10.1038/nmeth.2961.

(11) Dahlberg, P. D.; Saurabh, S.; Sartor, A. M.; Wang, J. R.; Mitchell, P. G.; Chiu, W.; Shapiro, L.; Moerner, W. E. Cryogenic single-molecule fluorescence annotations for electron tomography reveal in situ organization of key proteins in Caulobacter. Proceedings of the National Academy of Sciences of the United States of America 2020, 117 (25), 13937–13944. DOI: 10.1073/pnas.2001849117.

(12) Dahlberg, P. D.; Moerner, W. E. Cryogenic super-resolution fluorescence and electron microscopy correlated at the nanoscale. Annual Review of Physical Chemistry 2021, 72 (1), 253–278. DOI: 10.1146/annurev-physchem-090319-051546.

(13) DeRosier, D. J. Where in the cell is my protein? Quarterly Reviews of Biophysics 2021, 54, e9. DOI: 10.1017/S003358352100007X From Cambridge University Press Cambridge Core.

(14) Moerner, W. E. Microscopy beyond the diffraction limit using actively controlled single molecules. Journal of Microscopy 2012, 246 (3), 213–220. DOI: 10.1111/j.1365-2818.2012.03600.x.

(15) Möckl, L.; Moerner, W. E. Super-resolution microscopy with single molecules in biology and beyond - essentials, current trends, and future challenges. Journal of the American Chemical Society 2020, 142, 17828–17844. DOI: 10.1021/jacs.0c08178.

(16) Tuijtel, M. W.; Koster, A. J.; Jakobs, S.; Faas, F. G. A.; Sharp, T. H. Correlative cryo super-resolution light and electron microscopy on mammalian cells using fluorescent proteins. Scientific Reports 2019, 9 (1), 1369. DOI: 10.1038/s41598-018-37728-8 (acccessed 02/04).

(17) Mantovanelli, A. M. R.; Glushonkov, O.; Adam, V.; Wulffelé, J.; Thédié, D.; Byrdin, M.; Gregor, I.; Nevskyi, O.; Enderlein, J.; Bourgeois, D. Photophysical Studies at Cryogenic Temperature Reveal a Novel Photoswitching Mechanism of rsEGFP2. Journal of the American Chemical Society 2023, 145 (27), 14636–14646. DOI: 10.1021/jacs.3c01500.

(18) Dahlberg, P. D.; Sartor, A. M.; Wang, J. R.; Saurabh, S.; Shapiro, L.; Moerner, W. E. Identification of PAmKate as a red photoactivatable fluorescent protein for cryogenic super-resolution imaging. Journal of the American Chemical Society 2018, 140 (39), 12310–12313. DOI: 10.1021/jacs.8b05960.

(19) Lakowicz, J. R. Fluorophores. In Principles of Fluorescence Spectroscopy, 3 ed.; Springer, 2006; pp 63–67.

(20) Dubochet, J.; McDowall, A. W. VITRIFICATION OF PURE WATER FOR ELECTRON MICROSCOPY. Journal of microscopy 1981, 124 (3), 3–4. DOI: 10.1111/j.1365-2818.1981.tb02483.x; 14 10.1111/j.1365-2818.1981.tb02483.x (acccessed 12/01; 2020/05).

(21) Dahlberg, P. D.; Perez, D.; Hecksel, C. W.; Chiu, W.; Moerner, W. E. Metallic support films reduce optical heating in cryogenic correlative light and electron tomography. Journal of Structural Biology 2022, 214 (4), 107901. DOI: 10.1016/j.jsb.2022.107901.

(22) Voss, J. M.; Harder, O. F.; Olshin, P. K.; Drabbels, M.; Lorenz, U. J. Rapid melting and revitrification as an approach to microsecond time-resolved cryo-electron microscopy. Chemical Physics Letters 2021, 778, 138812. DOI: 10.1016/j.cplett.2021.138812.

(23) Sartor, A. M.; Dahlberg, P. D.; Perez, D.; Moerner, W. E. Characterization of mApple as a Red Fluorescent Protein for Cryogenic Single-Molecule Imaging with Turn-Off and Turn-On Active Control Mechanisms. The Journal of Physical Chemistry B 2023, 127 (12), 2690–2700. DOI: 10.1021/acs.jpcb.2c08995.

(24) Lower, S.; El-Sayed, M. The triplet state and molecular electronic processes in organic molecules. Chemical reviews 1966, 66 (2), 199–241.

(25) Campbell, R. E.; Tour, O.; Palmer, A. E.; Steinbach, P. A.; Baird, G. S.; Zacharias, D. A.; Tsien, R. Y. A monomeric red fluorescent protein. Proceedings of the National Academy of Sciences 2002, 99 (12), 7877–7882.

(26) Faro, A. R.; Adam, V.; Carpentier, P.; Darnault, C.; Bourgeois, D.; de Rosny, E. Low-temperature switching by photoinduced protonation in photochromic fluorescent proteins. Photochemical & Photobiological Sciences 2010, 9 (2), 254–262.

(27) Shcherbakova, D. M.; Verkhusha, V. V. Chromophore chemistry of fluorescent proteins controlled by light. Current Opinion in Chemical Biology 2014, 20, 60–68. DOI: 10.1016/j.cbpa.2014.04.010 (acccessed 06/01).

(28) Ausmees, N.; Kuhn, J. R.; Jacobs-Wagner, C. The bacterial cytoskeleton: an intermediate filament like function in cell shape. Cell 2003, 115 (6), 705–713. DOI: 10.1016/S0092-8674(03)00935-8.

(29) Briegel, A.; Dias, D. P.; Li, Z.; Jensen, R. B.; Frangakis, A. S.; Jensen, G. J. Multiple large filament bundles observed in Caulobacter crescentus by electron cryotomography. Mol. Microbiol. 2006, 62 (1), 5–14. DOI: 10.1111/j.1365-2958.2006.05355.x.

(30) Liu, Y.; van den Ent, F.; Löwe, J. Filament structure and subcellular organization of the bacterial intermediate filament–like protein crescentin. Proceedings of the National Academy of Sciences 2024, 121 (7), e2309984121. DOI: doi:10.1073/pnas.2309984121.

(31) Saurabh, S.; Perez, A. M.; Comerci, C. J.; Shapiro, L.; Moerner, W. E. Super-resolution Imaging of Live Bacteria Cells Using a Genetically Directed, Highly Photostable Fluoromodule. Journal of the American Chemical Society 2016, 138 (33), 10398–10401. DOI: 10.1021/jacs.6b05943 [doi].

(32) Ha, T.; Tinnefeld, P. Photophysics of Fluorescent Probes for Single-Molecule Biophysics and Super-Resolution Imaging. Annu. Rev. Phys. Chem. 2012, 63, 595–617.

(33) Acharya, A.; Bogdanov, A. M.; Grigorenko, B. L.; Bravaya, K. B.; Nemukhin, A. V.; Lukyanov, K. A.; Krylov, A. I. Photoinduced chemistry in fluorescent proteins: curse or blessing? Chemical Reviews 2017, 117 (2), 758–795. DOI: 10.1021/acs.chemrev.6b00238.

(34) English, D. S.; Harbron, E. J.; Barbara, P. F. Probing Photoinduced Intersystem Crossing by Two-Color, Double Resonance Single Molecule Spectroscopy. The Journal of Physical Chemistry A 2000, 104 (40), 9057–9061. DOI: 10.1021/jp001992y.

(35) Giske, A. CryoSTED microscopy-A new spectroscopic approach for improving the resolution of STED microscopy using low temperature. Ph. D., University of Heidelberg, 2007.

(36) Byrdin, M.; Duan, C.; Bourgeois, D.; Brettel, K. A long-lived triplet state is the entrance gateway to oxidative photochemistry in green fluorescent proteins. Journal of the American Chemical Society 2018, 140 (8), 2897–2905.

(37) Rane, L.; Wulffele, J.; Bourgeois, D.; Glushonkov, O.; Mantovanelli, A. M. R.; Zala, N.; Byrdin, M. Light-Induced Forward and Reverse Intersystem Crossing in Green Fluorescent Proteins at Cryogenic Temperatures. The Journal of Physical Chemistry B 2023, 127 (22), 5046–5054. DOI: 10.1021/acs.jpcb.3c02971.

(38) Vegh, R. B.; Bravaya, K. B.; Bloch, D. A.; Bommarius, A. S.; Tolbert, L. M.; Verkhovsky, M.; Krylov, A. I.; Solntsev, K. M. Chromophore photoreduction in red fluorescent proteins is responsible for bleaching and phototoxicity. The Journal of Physical Chemistry B 2014, 118 (17), 4527–4534. DOI: 10.1021/jp500919a.

(39) Ludvikova, L.; Simon, E.; Deygas, M.; Panier, T.; Plamont, M.-A.; Ollion, J.; Tebo, A.; Piel, M.; Jullien, L.; Robert, L.; et al. Near-infrared co-illumination of fluorescent proteins reduces photobleaching and phototoxicity. Nature Biotechnology 2024, 42 (6), 872–876. DOI: 10.1038/s41587-023-01893-7.

(40) Wulffele, J.; Thédié, D.; Glushonkov, O.; Bourgeois, D. mEos4b Photoconversion Efficiency Depends on Laser Illumination Conditions Used in PALM. The Journal of Physical Chemistry Letters 2022, 13 (22), 5075–5080. DOI: 10.1021/acs.jpclett.2c00933.

(41) Rego, E. H.; Shao, L.; Macklin, J. J.; Winoto, L.; Johansson, G. A.; Kamps-Hughes, N.; Davidson, M. W.; Gustafsson, M. G. L. Nonlinear structured-illumination microscopy with a photoswitchable protein reveals cellular structures at 50-nm resolution. Proc. Natl. Acad. Sci. U.S.A. 2012, 109, E135–E143.

(42) Cranfill, P. J.; Sell, B. R.; Baird, M. A.; Allen, J. R.; Lavagnino, Z.; de Gruiter, H. M.; Kremers, G. J.; Davidson, M. W.; Ustione, A.; Piston, D. W. Quantitative assessment of fluorescent proteins. Nat Methods 2016, 13 (7), 557–562. DOI: 10.1038/nmeth.3891 From NLM.

(43) Gunewardene, M. S.; Subach, F. V.; Gould, T. J.; Penoncello, G. P.; Gudheti, M. V.; Verkhusha, V. V.; Hess, S. T. Superresolution Imaging of Multiple Fluorescent Proteins with Highly Overlapping Emission Spectra in Living Cells. Biophys. J. 2011, 101 (6), 1522–1528. DOI: 10.1016/j.bpj.2011.07.049.

(44) Barentine, A. E. S.; Lin, Y.; Courvan, E. M.; Kidd, P.; Liu, M.; Balduf, L.; Phan, T.; Rivera-Molina, F.; Grace, M. R.; Marin, Z.; et al. An integrated platform for high-throughput nanoscopy. Nature Biotechnology 2023. DOI: 10.1038/s41587-023-01702-1.

(45) Kremer, J. R.; Mastronarde, D. N.; McIntosh, J. R. Computer Visualization of Three-Dimensional Image Data Using IMOD. Journal of structural biology 1996, 116 (1), 71–76. DOI: 10.1006/jsbi.1996.0013 (acccessed 01/01).

