## Supplementary Figures for "Exploring transient states of PAmKate to enable improved cryogenic single-molecule imaging"

Title

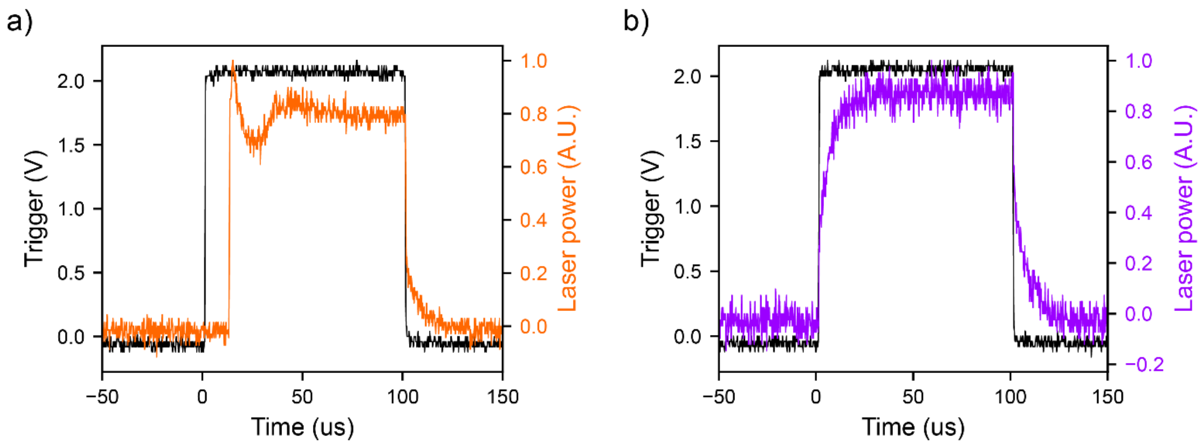


Figure S1: Laser pulse characterization of 594 nm (a) and 405 nm (b). The trigger signal sent to the digital modulation port of each laser is shown in black, and the response of the laser captured with a photodiode receiver (Newport 2001, 200 kHz bandwidth, gain factor 1) is shown in color. The 594 has a delay of about 10 µs followed by intensity fluctuation before stabilizing after about 50 µs.

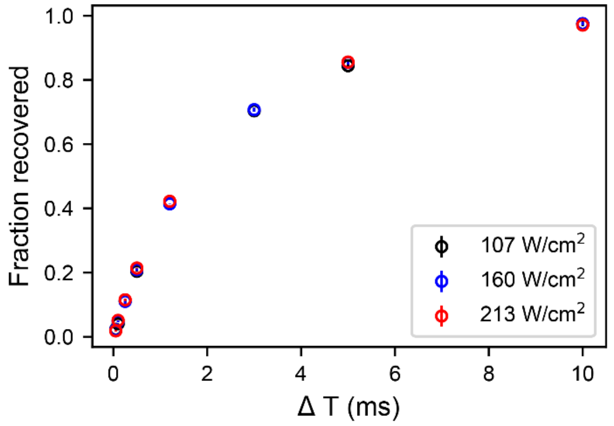


Figure S2: Recovery intensity dependence. Single exponential fit lifetimes were 2.3 ms, 2.3 ms, and 2.2 ms for 107, 160, and 213 W/cm$^2$ of 594 nm respectively. Standard deviation error bars are smaller than the points as drawn.

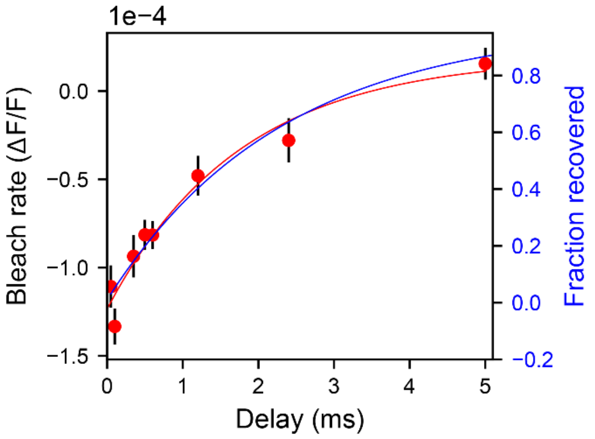


Figure S3: Comparing fits between transient bleaching (red) and recovery (blue). The bleach rate versus delay time for 594 + 405 nm (red points) and recovery fit (blue line) are the same as Figure 3c, and the red line is an exponential fit to the bleach rate versus delay time data.

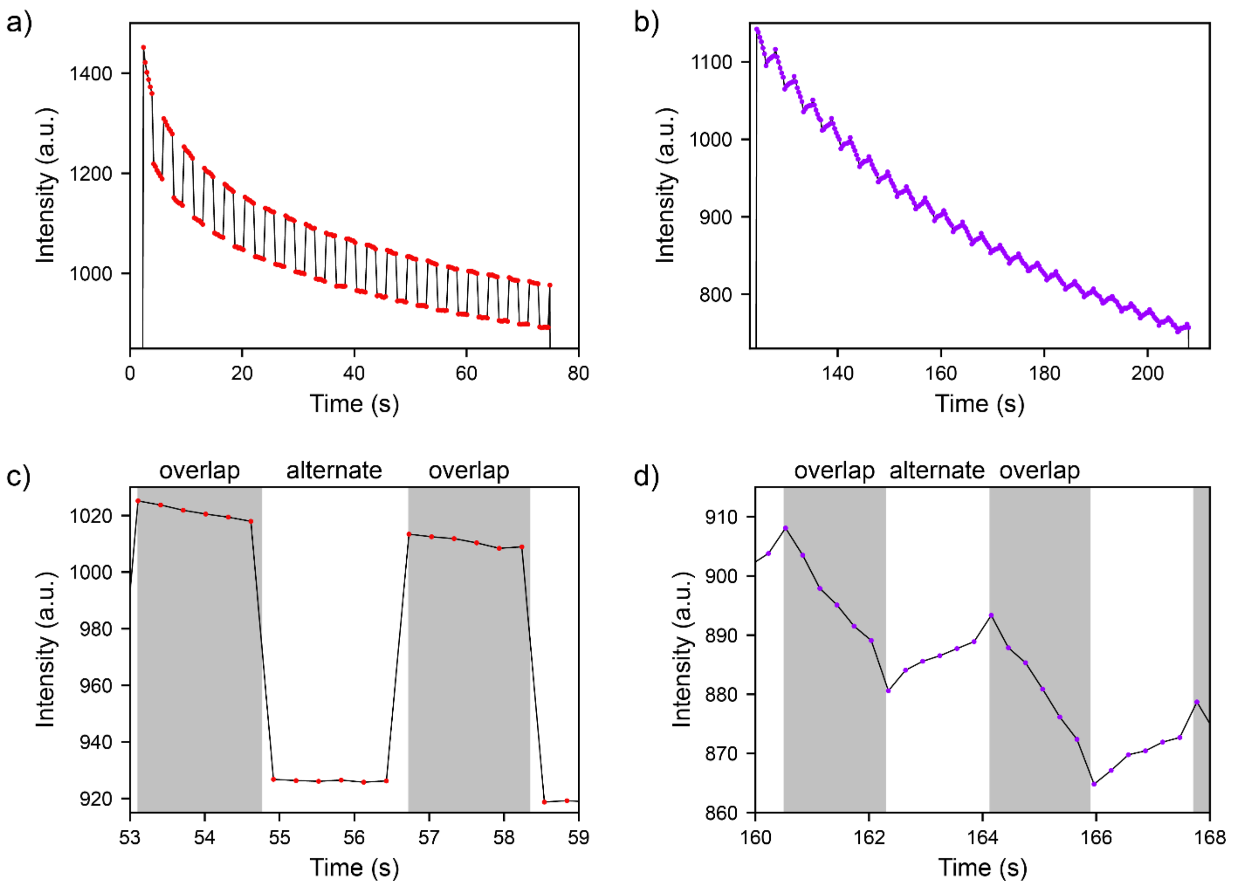


Figure S4: Action spectrum raw data for 594 + 720 nm (a, c) and 594 + 405 nm (b, d), taken from the same sample region. Following photoactivation with continuous 405 nm, the sample was illuminated with pulses of 594 and 720 nm, toggling between overlapping and alternating conditions. Then the routine was repeated with pulses of 594 and 405 nm. c) is a zoom in of a section of the data shown in a). Gray background marks when the pulses are overlapped, while white marks the alternating condition. When 594 and 720 nm are overlapped, the emission is brighter due to RISC or an analogous process, and the bleaching slope is slightly steeper. Likewise, d) is a zoom in from b). When 594 and 405 nm are overlapped, the fluorescence is slightly brighter than the alternating condition, and the bleaching has a steep negative slope. When the condition is switched to alternating, emission gets slightly dimmer, and the slope turns positive. This suggests the majority of the bleach by 405 nm of the transient state is irreversible, but some molecules are actually shelved reversibly, and can recover during the alternating condition. When the overlap condition resumes, the fluorescence is slightly brighter than the end of the previous overlap period, shown as a small magnitude brightening in Fig. 4b.

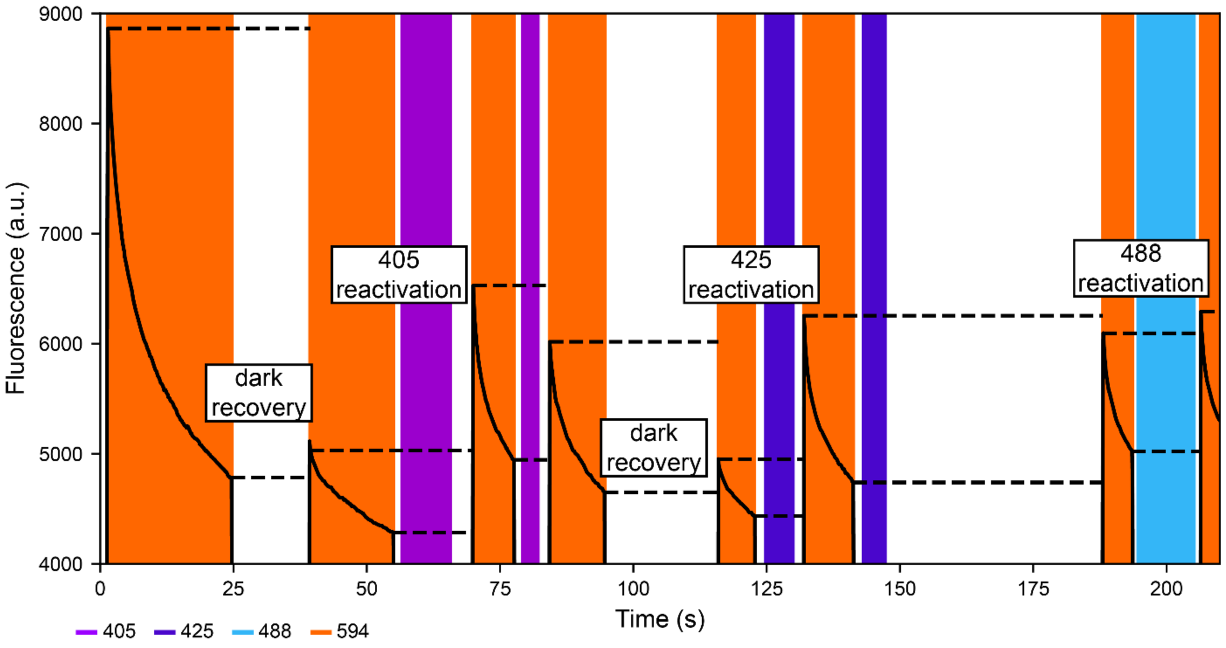


Figure S5: Photoswitching into and out of the long-lived dark state. Fluorescence from the sample is plotted with a black line, periods of exposure to various wavelengths are marked with colored backgrounds, and the initial and final brightness of each period of 594 exposure are marked with dashed lines. 594 exposure drives some molecules to permanent bleach and others to a long-lived dark state. Dark recovery results in a modest fluorescent increase. Exposure, even for short times, to 405, 425, and 488 photoswitch molecules back to the active state, resulting in a significant increase in brightness under 594 excitation. NIR (not shown here) is found to have no effect beyond dark recovery.

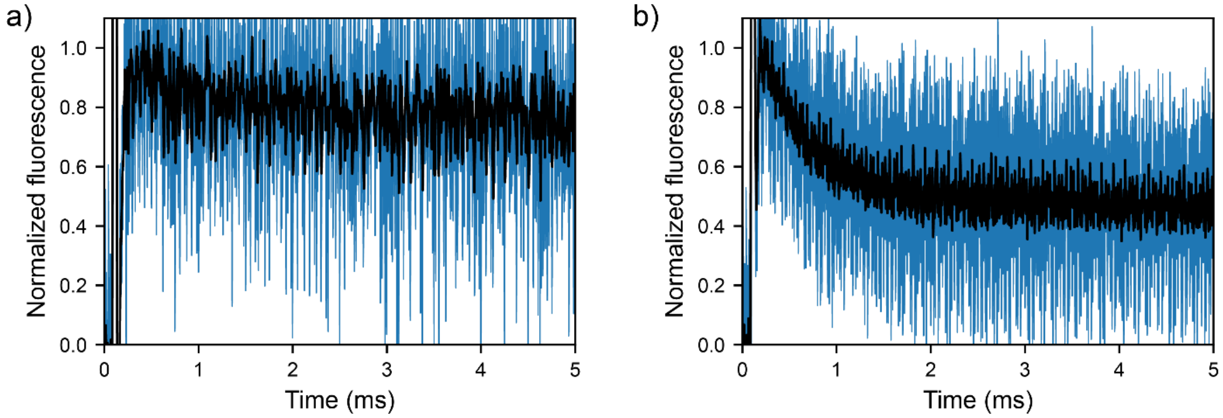


Figure S6: Shelving at RT (a) and CT (b) of spin coated thin film. To achieve adequate signal to noise in this thin sample, many scans over multiple regions are averaged to produce the traces in blue. A 10 μs sliding window average is shown in black. The spike at 0 ms is crosstalk from the trigger signal in the oscilloscope, but dissipates faster than the laser stabilizes. It is clear from this data that at RT the shelving is either not as efficient or much faster than at CT.

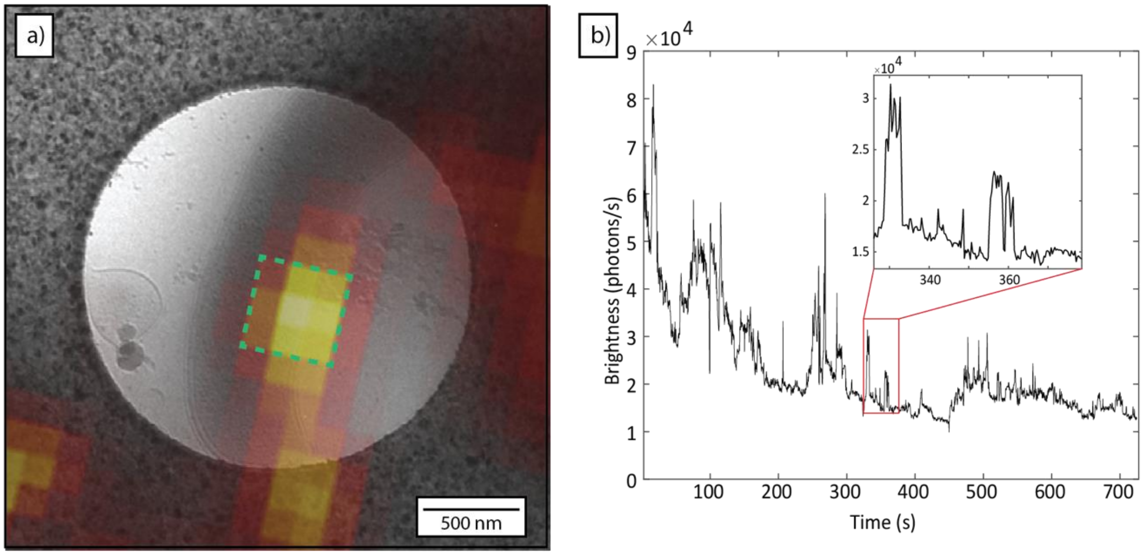


Figure S7: Fluorescence intensity trace for srCryoCLEM example. a) Diffraction-limited overlay of fluorescence (heat map) and low-magnification cryogenic electron micrograph of *C. crescentus* cell expressing PAmKate-CreS fusion construct. b) Integrated fluorescence brightness trace from pixel in green dashed box. Clear step-wise dynamics from single-molecule fluorescence are visible throughout the intensity trace. This indicates that even though 405 nm excitation was present throughout the acquisition the necessary sparsity of emitters was maintained.

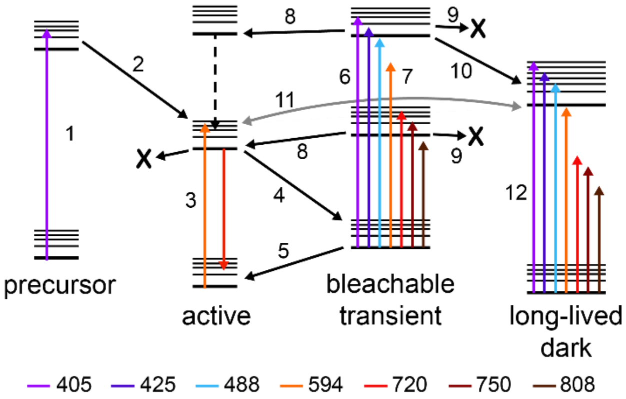


Figure S8: Annotated diagram of observed photophysical states. Same as Fig. 6 in the main text, with transitions annotated with numbers referring to Table S1.

| Arrow # | Description | Experimental evidence |
|---|---|---|
| 1 | Excitation of precursor | Previous work[1] |
| 2 | Photoactivation | Previous work[1] |
| 3 | Excitation/emission of fluorescence | Previous work[1] |
| 4 | Shelving to transient state | Fig. 2a |
| 5 | Recovery to active ground state | Fig. 2c |
| 6 | 405 excitation of transient state | Fig. 3c |
| 7 | Excitation of transient state with other $\lambda_2$ | Fig. 5 |
| 8 | Excited state return to active state | Fig. S4c/d |
| 9 | Bleach from excited transient state | Fig. 3c |
| 10 | Conversion of excited transient state to dark state | Fig. S4d |
| 11 | Photoconversion to and from dark state | Fig. S5 |
| 12 | Excitation of dark state | Fig. S5 |

Table S1: Annotation of transitions drawn in Fig. S8.

(1) Dahlberg, P. D.; Sartor, A. M.; Wang, J. R.; Saurabh, S.; Shapiro, L.; Moerner, W. E. Identification of PAmKate as a red photoactivatable fluorescent protein for cryogenic super-resolution imaging. *Journal of the American Chemical Society* **2018**, *140* (39), 12310-12313. DOI: 10.1021/jacs.8b05960.
